# Disruption of a Hedgehog-Foxf1-Rspo2 Signaling Axis Leads to Tracheomalacia and a Loss of Sox9+ Tracheal Chondrocytes

**DOI:** 10.1101/2020.07.11.198556

**Authors:** Talia Nasr, Praneet Chaturvedi, Kunal Agarwal, Jessica L. Kinney, Keziah Daniels, Stephen L. Trisno, Vladimir Ustiyan, John M. Shannon, James M. Wells, Debora Sinner, Vladimir V. Kalinichenko, Aaron M. Zorn

**Affiliations:** Center for Stem Cell and Organoid Medicine, Division of Developmental Biology, Perinatal Institute, Cincinnati Children’s Hospital Medical Center, Cincinnati, Ohio, 45229; Department of Pediatrics, University of Cincinnati College of Medicine, Cincinnati, Ohio, 45267; Division of Pulmonary Biology, Cincinnati Children’s Hospital Medical Center, Cincinnati, Ohio, 45229; Center for Lung Regenerative Medicine, Perinatal Institute, Cincinnati Children’s Hospital Medical Center, Cincinnati, Ohio, 45229

**Keywords:** Trachea, tracheomalacia, cartilage, Hedgehog

## Abstract

Congenital tracheomalacia, resulting from incomplete tracheal cartilage development, is a relatively common birth defect that severely impairs breathing in neonates. Mutations in the Hedgehog (HH) pathway and downstream Gli transcription factors are associated with tracheomalacia in patients and mouse models; however, the underlying molecular mechanisms are unclear. Using multiple *HH/Gli* mouse mutants including one that mimics Pallister-Hall Syndrome, we show that excessive Gli repressor activity prevents specification of tracheal chondrocytes. Lineage tracing experiments show that Sox9+ chondrocytes arise from HH-responsive splanchnic mesoderm in the fetal foregut that expresses the transcription factor Foxf1. Disrupted HH/Gli signaling results in 1) loss of Foxf1 which in turn is required to support Sox9+ chondrocyte progenitors and 2) a dramatic reduction in *Rspo2*, a secreted ligand that potentiates Wnt signaling known to be required for chondrogenesis. These results reveal a HH-Foxf1-Rspo2 signaling axis that governs tracheal cartilage development and informs the etiology of tracheomalacia.

**SUMMARY STATEMENT:** This work provides a mechanistic basis for tracheomalacia in patients with Hedgehog pathway mutations.

## INTRODUCTION

Impaired formation of the tracheal cartilage, or tracheomalacia, occurs in 1 in 2100 live births and can result in life-threatening airway collapse and impaired breathing (Boogaard et al., 2005; Kamran and Jennings, 2019). Current surgical treatment includes insertion of stents to keep the airway open, but these frequently lead to localized inflammation and multiple subsequent surgeries as the patients age (Fraga et al., 2016; Wallis et al., 2019). Generating biologically accurate replacement tissue from pluripotent stem cells is an aspirational strategy to improving patient care, but this requires a detailed understanding of both normal fetal tracheal development and the etiology of tracheomalacia (Fraga et al., 2016; Wallis et al., 2019).

Tracheal cartilage development in the mouse begins by embryonic day (E) 11.5 with expression of the transcription factor Sox9, a master regulator of chondrogenesis, in the ventral and lateral splanchnic mesenchyme surrounding the fetal trachea (Hines et al., 2013). Sox9+ cells do not condense around the dorsal side of the trachea, which forms the trachealis smooth muscle. Between E11.5 and E14.5, as the trachea continues to lengthen and grow, the Sox9+ presumptive chondrocytes organize into distinct, C-shaped rings separated by fibroelastic tissue along the anterior-posterior axis of the trachea (Kishimoto et al., 2018; Park et al., 2010). By E15.5, the chondrocytes differentiate into cartilage rings (Park et al., 2010). Hedgehog (HH) and Wnt signaling are critical for tracheal cartilage development in mice and mutations in these pathways have been associated with tracheomalacia in patients, but how these pathways interact to regulate tracheal chondrogenesis is unclear (Sinner et al., 2019).

The transcription factor Sox9 is required for the development of chondrocyte progenitors throughout the body (Lefebvre et al., 2019). Genetic deletion of *Wls*, encoding the cargo protein essential for Wnt ligand secretion from the tracheal epithelium leads to a loss of Sox9 expression in the tracheal mesenchyme and a failure in chondrocyte development, causing eventual tracheomalacia (Snowball et al., 2015). Mutations in a number of other Wnt ligands or receptors expressed in the fetal foregut including *Wnt4*, *Wnt5a*, *Wnt7b*, *Ror2*, and *Rspo2* also display deficits in cartilage development with varying extent of tracheomalacia (Bell et al., 2008; Caprioli et al., 2015; Kishimoto et al., 2018; Li et al., 2002).

Disruption in HH signaling can similarly result in tracheomalacia and loss of Sox9+ tracheal chondrocytes in mice (Litingtung et al., 1998; Miller et al., 2004; Motoyama et al., 1998; Park et al., 2010). The HH pathway regulates gene expression via zinc finger Gli transcription factors. In the absence of HH ligands, the HH receptor Smoothened is inhibited, leading to the proteolytical processing of Gli2 and Gli3 into isoforms that act as transcriptional repressors (GliR) (Briscoe and Therond, 2013). In the presence of HH, Smoothened is active, leading to the production of full-length Gli2 and Gli3 isoforms that activate target gene transcription (GliA). In general, Gli3 predominantly acts in the transcriptional repressor form, while Gli2 largely acts as a transcriptional activator (Litingtung et al., 2002; te Welscher et al., 2002; Vokes et al., 2008). Shh ligand is expressed in the developing foregut epithelium where it signals to the surrounding mesenchyme to regulate Gli activity (Ioannides et al., 2003). In *Shh*^−/−^ mutants, the primitive foregut tube fails to separate into distinct trachea and esophagus (Litingtung et al., 1998; Miller et al., 2004; Park et al., 2010). Cartilage never forms around the mutant foregut and there is a dramatic reduction in Sox9 expression and proliferation of the ventral foregut mesenchyme (Litingtung et al., 1998; Miller et al., 2004; Park et al., 2010). *Gli2*^−/−^;*Gli3*^+/−^ mouse embryos that have only one copy of Gli3 also exhibit tracheomalacia, whereas *Gli2*^+/−^;*Gli3*^−/−^ embryos, which lack Gli3 but have a single copy of Gli2, do not (Motoyama et al., 1998; Nasr et al., 2019). These data suggest that the balance of GliA to GliR is critical for normal tracheal development.

Indeed, Pallister-Hall Syndrome (PHS) [Online Mendelian Inheritance of Man (OMIM): 146510] patients have a heterozygous mutation in *GLI3* that leads to a truncated protein lacking the transcriptional activation domain. As a result, the mutant protein only has GLI3R transcriptional repression even in the presence of active HH signaling. PHS patients can exhibit multiple syndromic phenotypes and often present with laryngeal clefts and tracheomalacia (Bose et al., 2002; Johnston et al., 2005).

Thus, while both HH and Wnt are critical for tracheal development, how they functionally interact is unclear. Here, we use conditional *Smo*^*f/f*^ mouse mutants, which lack GliA, and *Gli3T*^*Flag*/+^ transgenic mice, which overexpress Gli3R, to show that imbalance of Gli activator and repressor activity disrupts specification of Sox9+ tracheal chondrocytes resulting in a tracheomalacia phenotype. We find that HH/Gli promotes the expression of Foxf1 in the ventral foregut mesenchyme, which in turn is required for Sox9 expression. Transcriptional profiling of *Foxg1Cre;Gli3T*^*Flag*/+^ foregut tissue reveals that in addition to loss of *Foxf1* and *Sox9*, there is a dramatic reduction in expression of *Rspo2*, a secreted ligand known to potentiate Wnt signaling which is required for cartilage development (Bell et al., 2008). In situ hybridization confirmed reduced expression of *Rspo2* as well as the Wnt response gene *Notum* in the ventral tracheal mesenchyme (Gerhardt et al., 2018). Re-analysis of published ChIP-seq data suggests that *Rspo2* is a direct transcriptional target of Foxf1. These data reveal a HH-Foxf1-Rspo2 axis where epithelial HH regulates Wnt signaling in the mesenchyme promoting the specification of Sox9+ tracheal chondrocytes.

## RESULTS

### Tracheal Chondrocytes Arise from the Splanchnic Foregut Mesoderm

In order to investigate the mechanisms of early tracheal chondrogenesis, we first performed lineage tracing experiments to confirm that the Sox9+ tracheal chondrocytes are derived from the lateral plate mesoderm and not the neural crest, which give rise to laryngeal cartilage (Tabler et al., 2017). For these experiments we crossed floxed *mT/mG* reporter mice to three different Cre lines: *Foxg1Cre* which recombines in the foregut mesendoderm beginning at E8.5; *Dermo1Cre* which recombines in the lateral plate mesoderm beginning at E10.5; or *Wnt1Cre* which recombines in the neural crest cells by E8.5 (Hebert and McConnell, 2000; Lewis et al., 2013; Li et al., 2008; Muzumdar et al., 2007; Ustiyan et al., 2018). At E13.5, the Foxg1Cre and Dermo1Cre expressing splanchnic mesoderm lineage traced Sox9+ tracheal chondrocytes, as well as the Foxf1+ mesenchyme and smooth muscle of the esophagus and dorsal trachea (Supplemental Figure 1A-C). In contrast, the Wnt1+ cells did not trace the tracheal chondrocytes or Foxf1+ smooth muscle (Supplemental Figure 1D), but, as previously reported, Wnt1-traced cells were only detected in laryngeal cartilage and neurons innervating the esophagus (data not shown) (Adachi et al., 2020; Tabler et al., 2017). This demonstrates that the laryngeal and tracheal cartilages have distinct origins with the latter arising from the lateral plate mesoderm.

### HH/Gli Imbalance Leads to Tracheomalacia

Pallister-Hall Syndrome patients, with a mutated copy of *GLI3* that leads to excessive *GLI3R*, frequently present with tracheomalacia (Bose et al., 2002; Johnston et al., 2005). To better understand how disrupted HH/Gli signaling results in tracheomalacia, we analyzed a series of conditional mouse mutants where we either deleted the HH receptor *Smo*, which effectively removes GliA, or we ectopically expressed *Gli3T*^*Flag*/+^, which like PHS patients has elevated Gli3R activity but preserved GliA function (Vokes et al., 2008). We also took advantage of the different times of *Foxg1Cre* and *Dermo1Cre* recombination to examine the temporal roles of HH/Gli activity.

At E15.5 all the *Gli3T*^*Flag*/+^ and *Smo*^*f/f*^ mutants showed varying degrees of tracheomalacia, with reduced cartilage development as indicated by Alcian Blue staining (Figure 1A). The early *Foxg1Cre* mutants were more severe than the later *Dermo1Cre* mutants. *Foxg1Cre;Smo*^*f/f*^ mutants had the most severe tracheomalacia, as well as tracheal stenosis and a hypoplastic foregut, while the later-acting *Dermo1Cre;Smo*^*f/f*^ mutant tracheas had relatively more cartilage than the other mutants. All mutants also showed varying losses of the dorsal trachealis muscle (Figure 1B) as well as some degrees of esophageal stenosis, supporting that HH/Gli signaling is also required for esophageal development (Jia et al., 2018; Litingtung et al., 1998).

**Figure 1:**
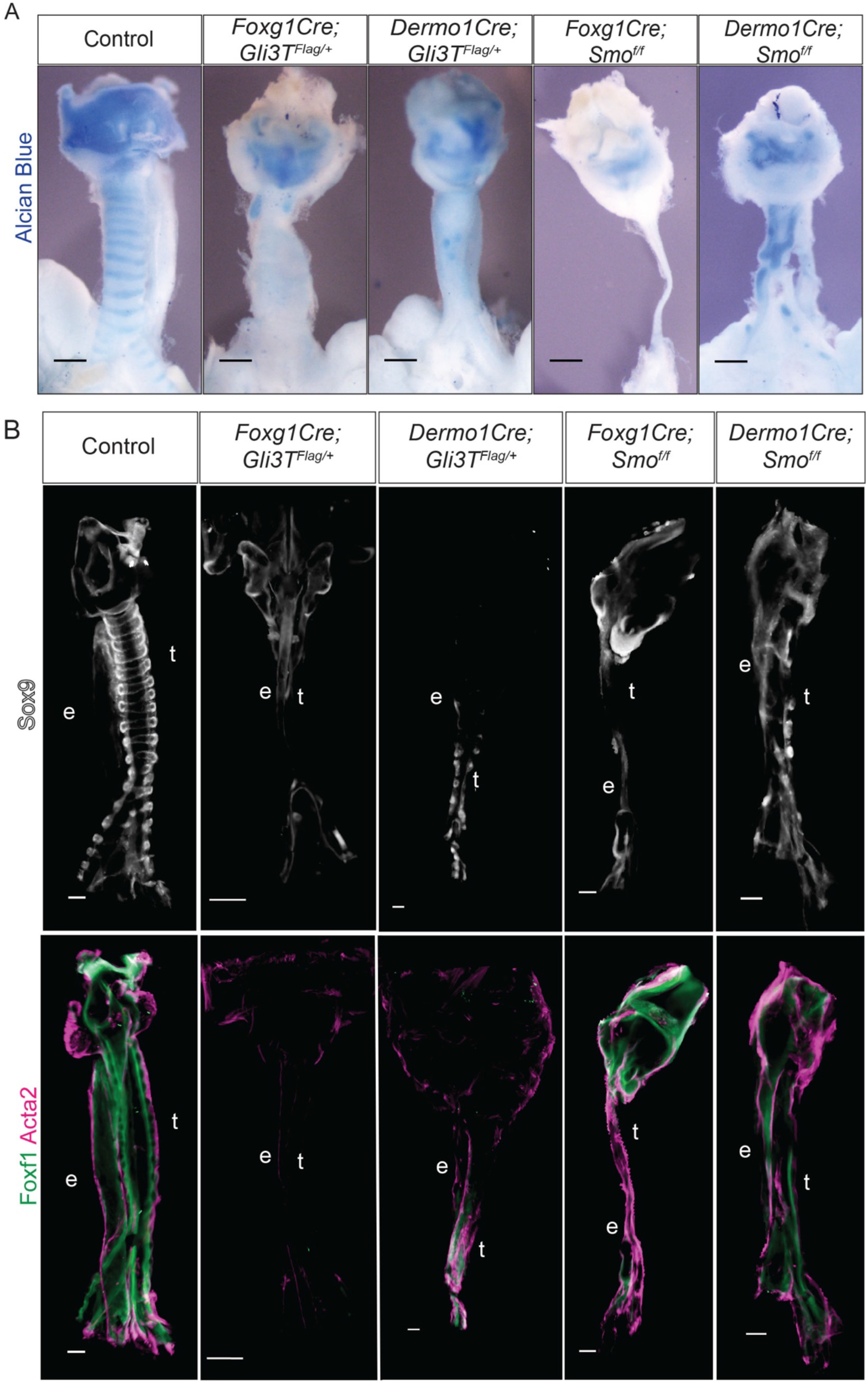
Imbalance in Gli Activity Leads to Tracheomalacia. A: Alcian Blue wholemounts of dissected E15.5 foreguts from control, *Foxg1Gli3T*^*Flag*/+^, *Dermo1Cre;Gli3T*^*Flag*/+^, *Foxg1Cre;Smo*^*f/f*^, and *Dermo1Cre;Smo*^*f/f*^ embryos. Earlier mutations generated using *Foxg1Cre* produced more severe tracheomalacia compared to *Dermo1Cre*-mediated deletions. N=3-5 embryos / genotype. B: Sox9, Foxf1, and Acta2 wholemount immunostaining of dissected E15.5 foreguts from control, *Foxg1Gli3T*^*Flag*/+^, *Dermo1Cre;Gli3T*^*Flag*/+^, *Foxg1Cre;Smo*^*f/f*^, and *Dermo1Cre;Smo*^*f/f*^ embryos. *Foxg1Cre* mutants display more significant reductions in Sox9 and Foxf1 compared to *Dermo1Cre* mutants, suggesting that impaired tracheal mesenchymal specification may contribute to tracheomalacia. N=3-5 embryos / genotype. All scale bars, 100 μm.

We next asked if Sox9+ tracheal chondrocytes were present at E15.5 but undifferentiated as a result of disrupting the HH/Gli pathway. However, all mutants showed reduced Sox9 levels that correlated with the level of Alcian Blue staining (Figure 1B), suggesting the loss of cartilage was not due primarily to a failure in differentiation, but rather due to a loss of Sox9+ chondrocytes. We also observed a reduction in Foxf1, a direct Gli target that is also required for foregut smooth muscle development (Hoffmann et al., 2014; Hoggatt et al., 2013; Ustiyan et al., 2018). Co-staining with Acta2 confirmed reduced smooth muscle differentiation, particularly in the *Gli3T* mutants (Figure 1B). Overall, more dramatic loss of Sox9+ tracheal chondrocytes and Foxf1+ muscle in *Foxg1Cre* mutants compared to *Dermo1Cre* mutants is consistent with HH/Gli signaling having a role in the foregut lateral plate mesoderm beginning at E8.5.

### Dynamic Foxf1 and Sox9 Localization during Tracheal Development

Since the phenotypes suggested an early disruption in chondrocyte development, we set out to better characterize the earliest expression of Sox9. Immunostaining showed that at E10.0, prior to separation of the foregut into distinct trachea and esophagus, the splanchnic mesoderm uniformly expressed Foxf1 with only rare interspersed Sox9+ cells, which based on our lineage tracing are likely to be neural crest-derived neurons (Figure 2A). Sox9 was first detected in the ventral-lateral mesoderm surrounding the trachea at E10.5 just after foregut separation, with the staining intensity and number of Sox9+ chondrocytes increasing by E11.5 (Figure 2A) (Hines et al., 2013). Initially Sox9 and Foxf1 were co-expressed, but as development proceeds, the upregulation of Sox9 in the ventral mesoderm was coincident with a downregulation of Foxf1. By E11.5, the Sox9 and Foxf1 expression domains were, for the most part, mutually exclusive with Foxf1 being restricted to the presumptive trachealis muscle, which previous studies have shown express Acta2 by this time, indicating a clear segregation of chondrocyte and smooth muscle lineages (Hines et al., 2013). Interestingly, Sox9/Foxf1 double positive cells persisted at the cartilage-smooth muscle boundary.

**Figure 2:**
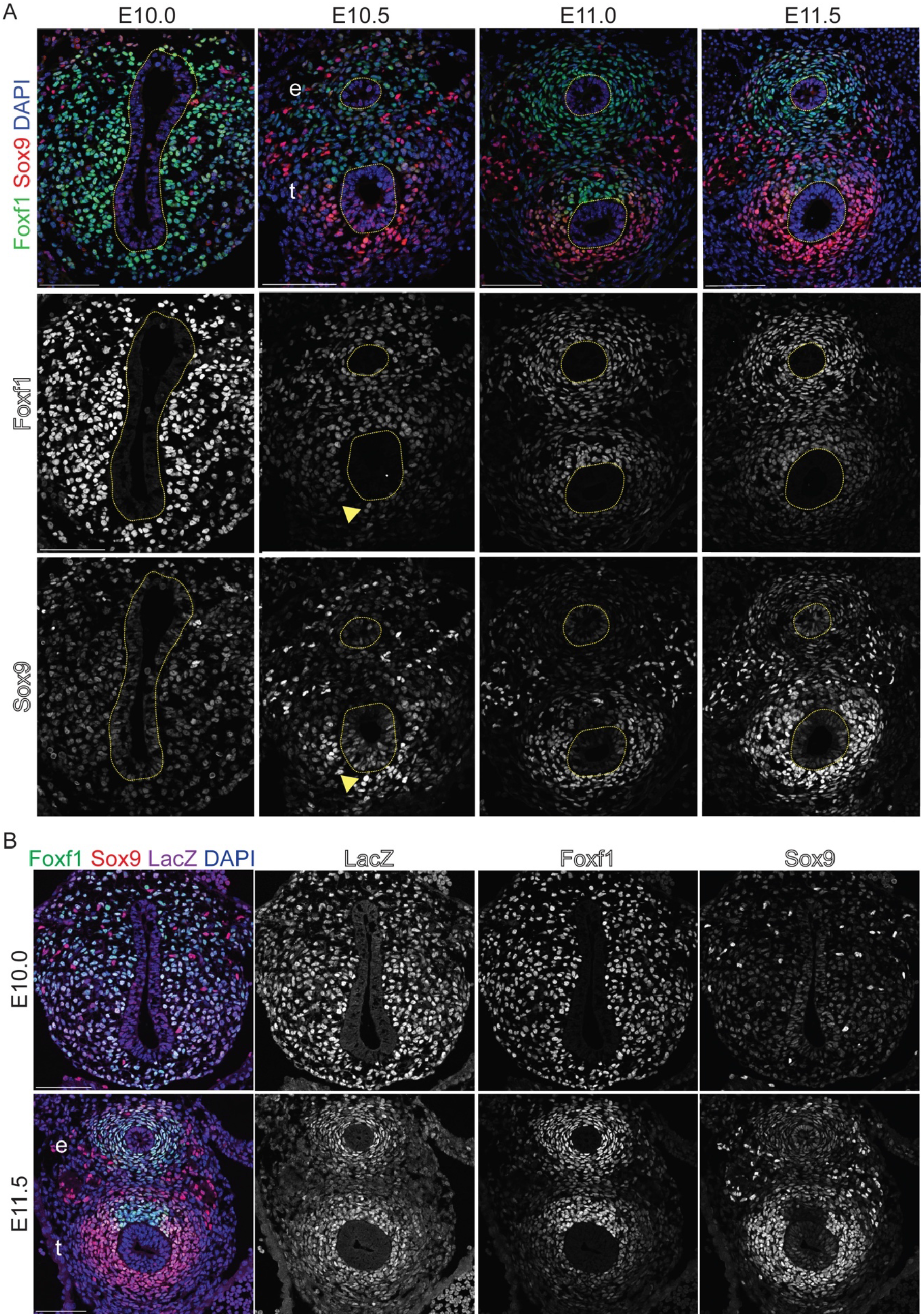
Dynamic Sox9 and Foxf1 Expression during Tracheal Chondrocyte Specification. A: Foxf1 and Sox9 immunostaining of control embryos between E10.0 – E11.5. Foxf1 is initially expressed throughout the lateral plate mesoderm, but is downregulated in the ventral tracheal mesoderm relative to the dorsal tracheal mesoderm by E11.5. Sox9 is only found in neural crest cells at E10.0, but is progressively localized in the ventral tracheal mesoderm between E10.5 and E11.5. Yellow arrowheads indicate Sox9-Foxf1 co-expressing cells in the E10.5 ventral tracheal mesoderm. N=3-5 embryos/stage B: Immunostaining of LacZ (ß-galactosidase), Foxf1, and Sox9 in E10.0 and E11.5 Gli1^LacZ/+^ embryos. Gli1 can be found throughout the lateral plate mesoderm surrounding the foregut at E10.0 and is enriched in the mesoderm immediately surrounding the esophageal and tracheal endoderm at E11.5. All scale bars, 100 μm. e=esophagus, t=trachea.

Since upregulation of Sox9 and downregulation of Foxf1 in the ventral tracheal mesenchyme follows tracheoesophageal separation, we next asked if this process was dependent on tracheoesophageal separation and/or epithelial identity. Nkx2-1 and Sox2 are transcription factors required for development of the tracheal and esophageal endoderm epithelial, respectively (Minoo et al., 1999; Que et al., 2007). *Nkx2-1*^−/−^ mutants have a single undivided foregut tube of esophageal character, whereas deletion of *Sox2* from the foregut results in an undivided foregut tube of respiratory character (Kuwahara et al., 2020; Que et al., 2009; Que et al., 2007; Teramoto et al., 2020; Trisno et al., 2018). A reanalysis of these mutants showed that both the *Sox2* and *Nkx2-1*-null embryos exhibited a downregulation of Foxf1 and upregulation of Sox9 in the ventral mesenchyme, although there were far fewer Sox9+ chondrocytes in the *Nkx2-1* mutant compared to the *Sox2* mutant or wildtype controls (Supplemental Figure 2A) (Que et al., 2009; Teramoto et al., 2020). Together, these data indicate that the emergence of tracheal chondrocytes with an upregulation of Sox9 and a downregulation of Foxf1 is influenced by the epithelial identity, but not dependent on tracheoesophageal separation.

### Hedgehog/Gli Activity is Required for Specification of the Tracheal Mesenchyme

Expression of Shh and Ihh ligands is known to be dynamic in the developing foregut. From E9.0-10.5, both *Shh* and *Ihh* have enriched expression in the ventral foregut epithelium, but after tracheoesophageal separation, *Shh* becomes downregulated in the trachea epithelium and upregulated in the esophageal epithelium (Ioannides et al., 2003; Rankin et al., 2016). Therefore, we examined whether dynamic HH signaling might account for the reciprocal Sox9-Foxf1 expression pattern, postulating that there might be an overall reduction of ventral HH activity that correlated with reduced Foxf1 and increased Sox9. We took advantage of *Gli1LacZ* reporter mice since Gli1 is a direct transcriptional target of HH-Gli2/3 signaling, enabling us to examine the overall impact of both Shh and Ihh activity (Briscoe and Therond, 2013). Contrary to our hypothesis, Gli1LacZ was uniformly expressed in the foregut mesoderm surrounding the gut tube at both at E10.0 and E11.5, co-localizing with the Foxf1+ smooth muscle and Sox9+ chondrocytes at E11.5 (Figure 2B). RNAScope *in situ* hybridization confirmed *Gli1* and *Shh* expression patterns (Supplemental Figure 2B) (Ioannides et al., 2003). We also observed uniform Smo expression around the tracheal endoderm, supporting that HH/Gli signaling is still present in the ventral tracheal mesoderm at E11.5 (Supplemental Figure 2B). Thus, the splanchnic mesoderm is actively responding to epithelial HH signaling during Sox9+ chondrocyte specification.

Next, we performed Foxf1 and Sox9 immunostaining on *Gli3T*^*Flag*/+^ and *Smo*^*f/f*^ mutants at E11.5 to examine the initial defects in tracheal chondrogenesis (Figure 3). The *Foxg1Cre;Smo*^*f/f*^ and *Foxg1Cre;Gli3T*^*Flag*/+^ mutants had reduced number of Sox9+ chondrocytes and also exhibited reduced ventral Foxf1 expression compared to controls (Figures 3A-D). *Dermo1Cre* mutants appeared to be mostly unchanged in both overall tracheal mesoderm cell number and lineage-specific populations, although they did exhibit a trend of reduced Foxf1 expression levels (Figure 3A-C). Since we observe loss of Sox9 and Foxf1 at E15.5 in both *Dermo1Cre* mutants, this suggests a continuing role for HH/Gli signaling in maintaining Foxf1 and Sox9 expression and promoting tracheal chondrogenesis.

**Figure 3:**
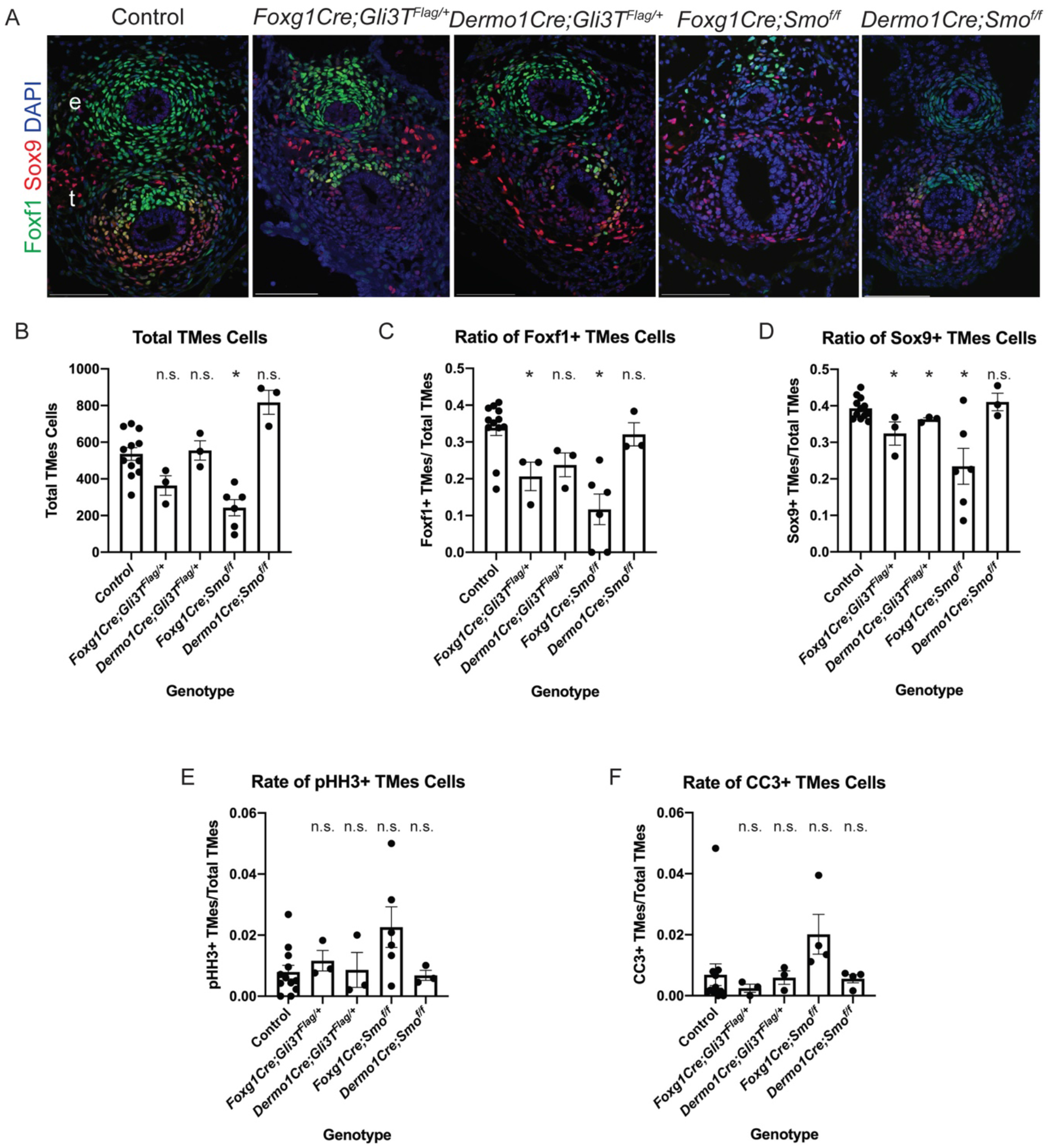
Hedgehog/Gli Signaling Supports Tracheal Mesenchymal Specification. A: Transverse sections of the E11.5 foregut in control, *Foxg1Cre;Gli3T*^*Flag*/+^, *Dermo1Cre;Gli3T*^*Flag*/+^, *Foxg1Cre;Smo*^*f/f*^, and *Dermo1Cre;Smo*^*f/f*^ embryos. Foxf1 and Sox9 immunostaining shows that *Foxg1Cre* mutants have fewer mesenchymal cells, along with fewer Foxf1+ and Sox9+ cells compared to *Dermo1Cre* mutants and control embryos. However, Sox9+ enteric neural crest cells are still present. N=3-5 embryos / genotype. All scale bars, 100 μm. e=esophagus, t=trachea. B: Quantification of the total number of cells in the tracheal mesoderm (TMes) in E11.5 *Gli3T*^*Flag*/+^ and *Smo*^*f/f*^ mutants. *Foxg1Cre;Smo*^*f/f*^ and *Foxg1Cre;Gli3T*^*Flag*/+^ mutants exhibit fewer tracheal mesoderm cells compared to controls, indicating that early disruptions in HH/Gli signaling lead to a reduction in tracheal mesenchyme. C: Quantification of the ratio of Foxf1+ or Sox9+ cells in the tracheal mesoderm (TMes) in E11.5 *Gli3T*^*Flag*/+^ and *Smo*^*f/f*^ mutants. *Foxg1Cre;Smo*^*f/f*^ and *Foxg1Cre;Gli3T*^*Flag*/+^ mutants in particular demonstrate relative reduction in Foxf1+ and Sox9+ cells. *Dermo1Cre;Gli3T*^*Flag*/+^ mutants also show reduction in Sox9+ and Foxf1+ cells while *Dermo1Cre;Smo*^*ff*^ do not, suggesting that earlier *Foxg1Cre*-mediated recombination results in more severe disruptions of tracheal mesenchyme patterning. D: Quantification of the cell proliferation rate, or mitotic index, in E11.5 *Gli3T*^*Flag*/+^ and *Smo*^*f/f*^ tracheal mesoderm (TMes) cells using phospho-histone H3 (pHH3) immunostaining to mark proliferating cells. Overall, disruptions in cell proliferation appear to be minimal in *Gli3T*^*Flag*/+^ and *Smo*^*f/f*^ mutants, indicating that reduced cell proliferation at E11.5 cannot explain the observed tracheomalacia phenotypes. E: Quantification of tracheal mesoderm (TMes) cell death rates in *Gli3T*^*Flag*/+^ and *Smo*^*f/f*^ mutants using cleaved caspase-3 (CC3) immunostaining to mark apoptotic cells. Disruption of HH/Gli signaling does not appear to affect the rate of cell death in the developing trachea, indicating that increased apoptosis at E11.5 cannot account for tracheomalacia seen in these mutants. All quantification shown as mean ± standard error of the mean, with p<0.05 as calculated by a two-sided Student’s t-test with unequal variance.

Since HH signaling is known to maintain cell proliferation and survival in many contexts, we assessed whether this might contribute to the tracheomalacia phenotype in *Smo*^*f/f*^ and *Gli3T*^*Flag*/+^ mutants (Bohnenpoll et al., 2017; Li et al., 2004). At E11.5 there were no statistically significant changes in either tracheal mesodermal cell proliferation or apoptosis in any of the mutants as determined by quantification of phospho-Histone H3 or cleaved caspase-3 immunostaining, respectively (Figures 3E-F). However, previous studies have demonstrated that HH/Gli does indeed promote splanchnic mesoderm proliferation and survival from E8.5 to E9.5, which likely explains the reduced cell number in *Foxg1Cre* mutants, which recombine starting at E8.5 (Figure 3B) (Li et al., 2008; Rankin et al., 2016). Overall, however, the reduced cell numbers in *Foxg1Cre* mutant is not sufficient to explain the loss of Foxf1 and Sox9. Together the results of Figures 1-3 indicate that HH/Gli is required between E8.5-10.5 to maintain Foxf1 and specify Sox9+ chondrocytes. Then, between E10.5 and 15.5, prolonged HH signaling is essential to maintain cartilage and smooth muscle development.

### Foxf1 Is Required for Development of Sox9+ Chondrocytes

Previous work has shown that loss of one *Foxf1* allele as well as conditional loss of *Foxf1* leads to impaired tracheal and esophageal development (Mahlapuu et al., 2001; Ustiyan et al., 2018). The reciprocal expression pattern of Foxf1 and Sox9 and the fact that both are reduced in the *Smo*^*f/f*^ and *Gli3T*^*Flag*/+^ mutants suggest that Foxf1 may initially be required for the development of Sox9+ progenitors, but that Sox9 and Foxf1 might then antagonize each other’s expression. To test this, we conditionally deleted *Foxf1* using both *Foxg1Cre* and *Dermo1Cre*. *Foxg1Cre;Foxf1*^*f/f*^ mutants exhibited a large reduction in Sox9+ ventral foregut mesoderm cells (Figure 4A), consistent with the small cartilaginous nodules previously observed in *Foxf1*^+/−^ tracheas (Mahlapuu et al., 2001). *Dermo1Cre;Foxf1*^*f/f*^ mutants also exhibited fewer Sox9+ cells compared to controls (Figure 4A). Quantification of cell numbers revealed that at E11.5, *Foxf1* mutants had a general trend of around 200 fewer tracheal mesoderm cells compared to controls, while only the *Foxg1Cre;Foxf1*^*f/f*^ mutants had significantly fewer Sox9+ cells in the tracheal mesoderm (Figures 4C-E). Interestingly, we observed that a number of the remaining Sox9+ cells also expressed Foxf1, suggesting that they escaped Cre recombination (Figure 4A, arrowhead). The fact that Sox9 was not upregulated in the *Dermo1Cre;Foxf1*^*f/f*^ mutants indicates that Foxf1 does not repress Sox9, which was one possibility suggested by their reciprocal expression patterns. We also found that the *Foxg1Cre;Foxf1*^*f/f*^ mutant foregut failed to separate into a distinct trachea and esophagus (Figure 4A), consistent with our recent work suggesting that the early lateral plate mesoderm is required for morphogenesis (Nasr et al., 2019).

**Figure 4:**
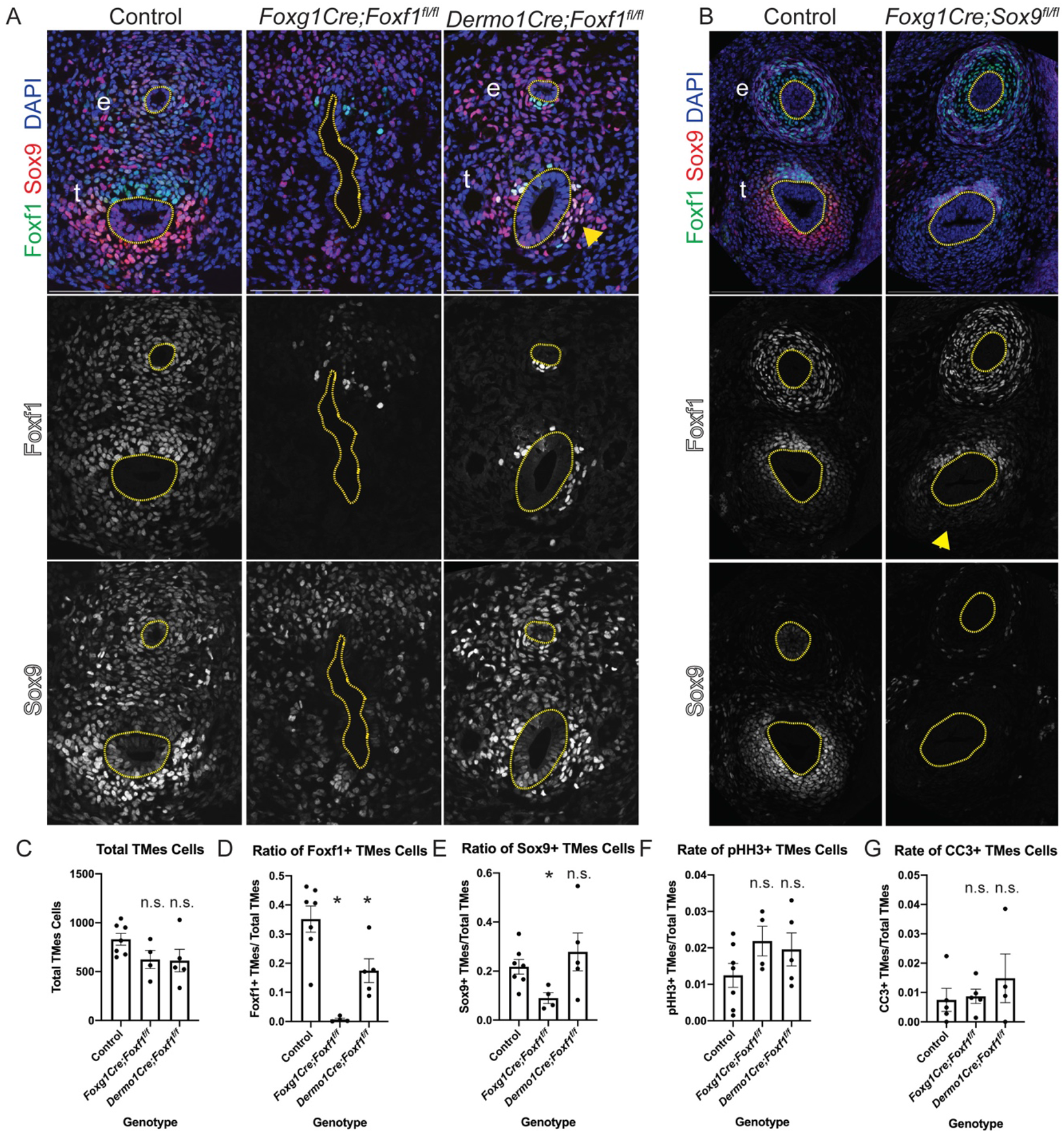
Foxf1 Is Required for Tracheal Sox9 Expression. A: Immunostaining of Foxf1 and Sox9 in E11.5 control, *Foxg1Cre;Foxf1*^*f/f*^, and *Dermo1Cre;Foxf1*^*f/f*^ mutants. *Foxg1Cre* mutants show a large reduction in both Foxf1 and Sox9 compared to controls. *Dermo1Cre* mutants have some Sox9+ cells in the ventral trachea that co-localize with or are immediately adjacent to Foxf1+ cells that escaped Cre recombination (yellow arrowhead). B: Foxf1 and Sox9 immunostaining in E13.5 control and *Foxg1Cre;Sox9*^*f/f*^ mutants. Compared to controls, *Foxg1Cre;Sox9*^*f/f*^ mutants have absent Sox9 expression in the ventral trachea but preserve downregulation of Foxf1 (yellow arrowhead). C: Quantification of the total number of tracheal mesoderm (TMes) cells in E11.5 *Foxf1*^*f/f*^ mutants suggests that loss of Foxf1 in the tracheal mesoderm beginning at E8.5 adversely affects tracheal mesoderm growth. D: Quantification of the ratio of Foxf1+ or Sox9+ cells in the tracheal mesoderm (TMes) in E11.5 *Foxf1*^*f/f*^ mutants indicates a relative reduction in both Foxf1+ and Sox9+ cells in *Foxg1Cre;Foxf1*^*f/f*^ mutants. E: Tracheal mesenchyme (TMes) mitotic indices in control, *Foxg1Cre;Foxf1*^*f/f*^, and *Dermo1Cre;Foxf1*^*f/f*^ embryos as determined by co-localization with pHH3 immunostaining. F: Quantification of apoptosis in tracheal mesenchymal (TMes) cells in control, *Foxg1Cre;Foxf1*^*f/f*^, and *Dermo1Cre;Foxf1*^*f/f*^ embryos as determined by CC3 immunostaining. All scale bars, 100 μm. e=esophagus, t=trachea. All quantification shown as mean ± standard error of the mean, with p<0.05 as calculated by a two-sided Student’s t-test with unequal variance.

We next asked if Sox9 might suppress the ventral expression of Foxf1 as chondrocytes emerge. Examination of E13.5 *Foxg1Cre;Sox9*^*f/f*^ mutants revealed that, loss of Sox9 had no impact on the ventral downregulation of Foxf1 (Figure 4B). This suggests that Sox9 and its downstream targets in the tracheal mesoderm are not responsible for the ventral reduction in Foxf1 expression. Thus, Foxf1 and Sox9 do not repress each other’s expression during tracheal chondrogenesis.

Since previous work showed that few *Dermo1Cre;Foxf1*^*f/f*^ mutants survive beyond E16.5 due to impaired growth and survival of the cardiovascular and pulmonary mesenchyme (Ustiyan et al., 2018), we examined cell proliferation and apoptosis. However, at E11.5 we did not detect any significant differences in cell proliferation or cell apoptosis in either *Foxg1Cre;Foxf1*^*f/f*^ or *Dermo1Cre;Foxf1*^*f/f*^ compared to controls (Figures 4F and 4G). As neither changes in cell proliferation nor cell death can explain the relative reduction in Sox9+ cells in *Foxg1Cre;Foxf1*^*f/f*^ mutants, we conclude that Foxf1 is required for initial specification of Sox9+ tracheal chondrocyte.

### Gli-Foxf1 Signaling Regulates Expression of *Rspo2*, a Known Wnt Modulator of Tracheal Chondrogenesis

Previous studies have shown that both *Foxf1* and *Sox9* are direct transcriptional targets of HH/Gli in different cellular contexts (Bien-Willner et al., 2007; Hoffmann et al., 2014; Madison et al., 2009; Tan et al., 2018). In order to discover additional Gli-regulated genes that might mediate tracheal chondrogenesis, we performed RNA sequencing on E10.5 foreguts and E11.5 tracheas dissected from control and *Foxg1Cre;Gli3T*^*Flag*/+^ embryos. Differential expression analysis (Log_2_ Fold Change >|1|, p<0.05) identified 708 transcripts (70 reduced and 638 increased) with altered expression in mutants at E10.5 and 738 Gli-regulated transcripts (352 reduced and 386 increased) at E11.5 (Figures 5A and B, Supplemental Table 1). Of these, 144 genes were differentially expressed at both E10.5 and E11.5. The reduced expression of *Hhip*, a direct HH target gene, was consistent with Gli3T repressive activity (Beachy et al., 2010). Gene ontology (GO) enrichment analysis of the downregulated genes identified epithelial tube morphogenesis and respiratory system development, consistent with HH signaling being required for foregut organogenesis, whereas upregulated genes were associated with cell signaling and muscle development, indicative of the relative increase in muscle progenitors in the absence of Sox9+ chondrocytes (Supplemental Figure S3A-B).

**Figure 5:**
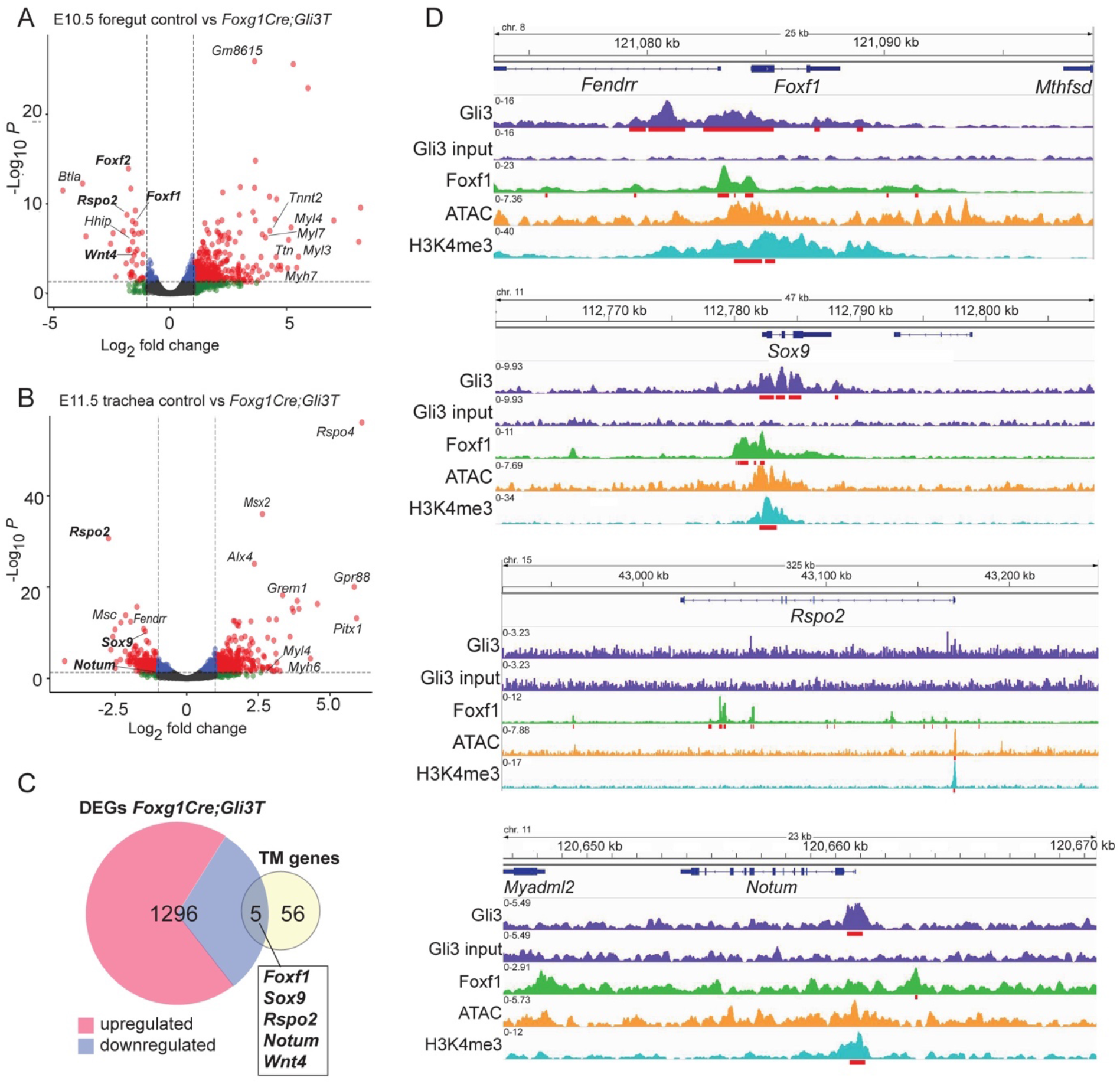
Gli3 Regulates Expression of Wnt Pathway Components. A: Volcano plot of differentially expressed transcripts in E10.5 control versus *Foxg1Gli3T*^*Flag*/+^ foreguts as determined by Log_2_FC ≥|1|, p<0.05. B: Volcano plot of differentially expressed transcripts in E11.5 control versus *Foxg1Gli3T*^*Flag*/+^ tracheas as determined by Log_2_FC ≥|1|, p<0.05. C: Venn diagram intersecting genes differentially expressed in *Foxg1Cre;Gli3T*^*Flag*/+^ mutants with genes known to be involved in human and mouse tracheal chondrogenesis (Shefchek et al., 2020; Sinner et al., 2019). TM = tracheomalacia-associated genes. D: Genome browser views of Gli3-3xFlag (GSE133710), Foxf1 (GSE77159), and H3K4me3 (GSE119885) ChiP-seq data, as well as ATAC-seq data (GSE119885) on *Foxf1*, *Sox9*, *Rspo2*, and *Notum* loci. Gli3 and Foxf1 bind the *Foxf1*, *Sox9*, and *Notum* loci, but only Foxf1 shows direct binding in the *Rspo2* locus along with significant ATAC and H3K4me3 peaks suggesting active transcription. Statistically significant ChIP peaks are underlined in red.

We next intersected the Gli3R-regulated transcripts with a manually curated list of 61 genes implicated in tracheal chondrogenesis and/or tracheomalacia in mice or humans (Supplemental Table 2). These were identified from a review of the literature (Sinner et al., 2019) (Supplemental Figure 3C) and by searching the Monarchinitiative.org, an online knowledgebase that aggregates human disease and animal model genotype-phenotypes associations (Shefchek et al., 2020). This intersection revealed five genes, all of which were downregulated in Gli3T transgenic embryos (Figure 5C), consistent with them being direct targets of Gli3 repressor activity. In addition to *Sox9* and *Foxf1*, this identified *Rspo2, Wnt4*, and *Notum*, all key regulators of the canonical Wnt pathway (Figure 5A-C and Supplemental Figure 3D-E) and all of which exhibit impaired tracheal chondrogenesis when mutated in mice (Bell et al., 2008; Caprioli et al., 2015; Gerhardt et al., 2018). Focusing on the Wnt pathway, we additionally observed reduced expression of *Wnt11* (Supplemental Figure 3C), whose role in tracheal development has not yet been identified but is known to support Sox9+ chondrocyte maturation in other tissues (Liu et al., 2014; Tada and Smith, 2000). *Rspo2*, a secreted protein that interacts with Lgr4/5/6 and Lrp6 receptor complexes to potentiate Wnt/β-catenin signaling, was one of the most downregulated transcripts at both E10.5 and 11.5 (−1.86 and −2.73 Log_2_FC, respectively) (Bell et al., 2008; Carmon et al., 2011; de Lau et al., 2011; Gong et al., 2012; Kazanskaya et al., 2004; Kim et al., 2008; Lebensohn and Rohatgi, 2018; Ruffner et al., 2012). *Wnt4* was modestly downregulated in the E10.5 foregut (−1.54 Log_2_FC), whereas Notum, a known Wnt target gene and Wnt feedback inhibitor was reduced about two-fold in the E11.5 Gli3T trachea (Figure 5B-C, Supplemental Figure 3C) (Gerhardt et al., 2018). These data demonstrate that HH/Gli signaling transcriptionally regulate components of the canonical Wnt pathway, which are known to activate Sox9 expression in the tracheal mesenchyme (Snowball et al., 2015).

We next examined published ChIP-seq data to examine whether *Rspo2*, *Notum*, *Wnt4*, and *Wnt11* were likely to be direct target genes of Gli and Foxf1 transcription factors. We used previously published Gli3-3xFlag ChIP from E10.5 limb buds and Foxf1 ChIP from E18.5 lungs (Dharmadhikari et al., 2016; Lex et al., 2020); these datasets are the most similar to tracheal chondrocytes currently available. We also examined previously published ATAC-seq and H3K4me3 ChIP-seq performed in the E9.5 cardiopulmonary foregut progenitors to help identify active promoter and enhancer regions (Steimle et al., 2018). Examination of genome browsers showed that Gli3 was bound to *Foxf1* and *Sox9* promoter regions that overlapped with H3K4me3 peaks (Figure 5D), consistent with previous reports that they are direct HH/Gli targets (Hoffmann et al., 2014; Tan et al., 2018; Vokes et al., 2008). Gli3 binding regions were also detected on putative regulatory elements of *Notum* and *Wnt 11* but not on the *Rspo2* or *Wnt4* loci (Figure 5D and Supplemental Figure 3F), suggesting that *Rspo2* and *Wnt4* might be indirectly regulated by Gli. Indeed, the Foxf1 ChIP-seq data showed that Foxf1 associated with intronic regions of *Rspo2*; the *Sox9*, *Notum*, *Wnt4* and *Wnt11* loci; as well as *Foxf1*’s own promoter (Figure 5D and Supplemental Figure 3F). The binding of Foxf1 to the *Sox9* promoter is consistent with direct regulation and the loss of Sox9 expression in *Foxg1Cre;Foxf1*^*f/f*^ mutants (Figures 4A and 4D). In addition, previous analysis of the same Foxf1 ChIP data in the embryonic lung reported Foxf1 chromatin binding to the *Wnt11* locus (Ustiyan et al., 2018). Although the ChIP data are not from the developing trachea, together with the RNA-seq, these data are consistent with Gli3-Foxf1 acting in a regulatory network to promote expression of *Sox9, Foxf1, Rspo2, Notum*, *Wnt4*, and *Wnt11* in the presumptive chondrocytes.

### Wnt Signaling Is Disrupted in *Gli3R* and *Foxf1* Mutants

We performed RNAscope *in situ* hybridization on E11.5 embryos to validate the RNA-seq analysis and examine which cell populations exhibited a change in *Rspo2*, *Notum* and *Wnt4* expression. In controls, *Rspo2* and *Notum* were strongly expressed in the ventral tracheal mesoderm, whereas *Wnt4* was weakly expressed in the mesoderm surrounding the esophagus and trachea as well as the epithelium (Figures 6A and 6B, Supplemental Figure 3E), all consistent with previous publications (Bell et al., 2008; Caprioli et al., 2015; Gerhardt et al., 2018). In both *Foxg1Cre;Gli3T*^*Flag*/+^ and *Foxg1Cre;Foxf1*^*f/f*^ mutants, *Rspo2* and *Notum* were dramatically downregulated, while there was only a modest reduction of *Wnt4* levels in half the embryos (Figures 6A and 6B, Supplemental Figure 3E). Since Notum is a direct Wnt target gene and is required for tracheal chondrogenesis, this suggests that Wnt signaling is disrupted in the presumptive chondrocytes of *Foxg1Cre;Gli3T*^*Flag*/+^ embryos (Gerhardt et al., 2018). Together, these data demonstrate that HH/Gli regulate a Foxf1-Wnt pathway to promote Wnt signaling and tracheal chondrogenesis.

**Figure 6:**
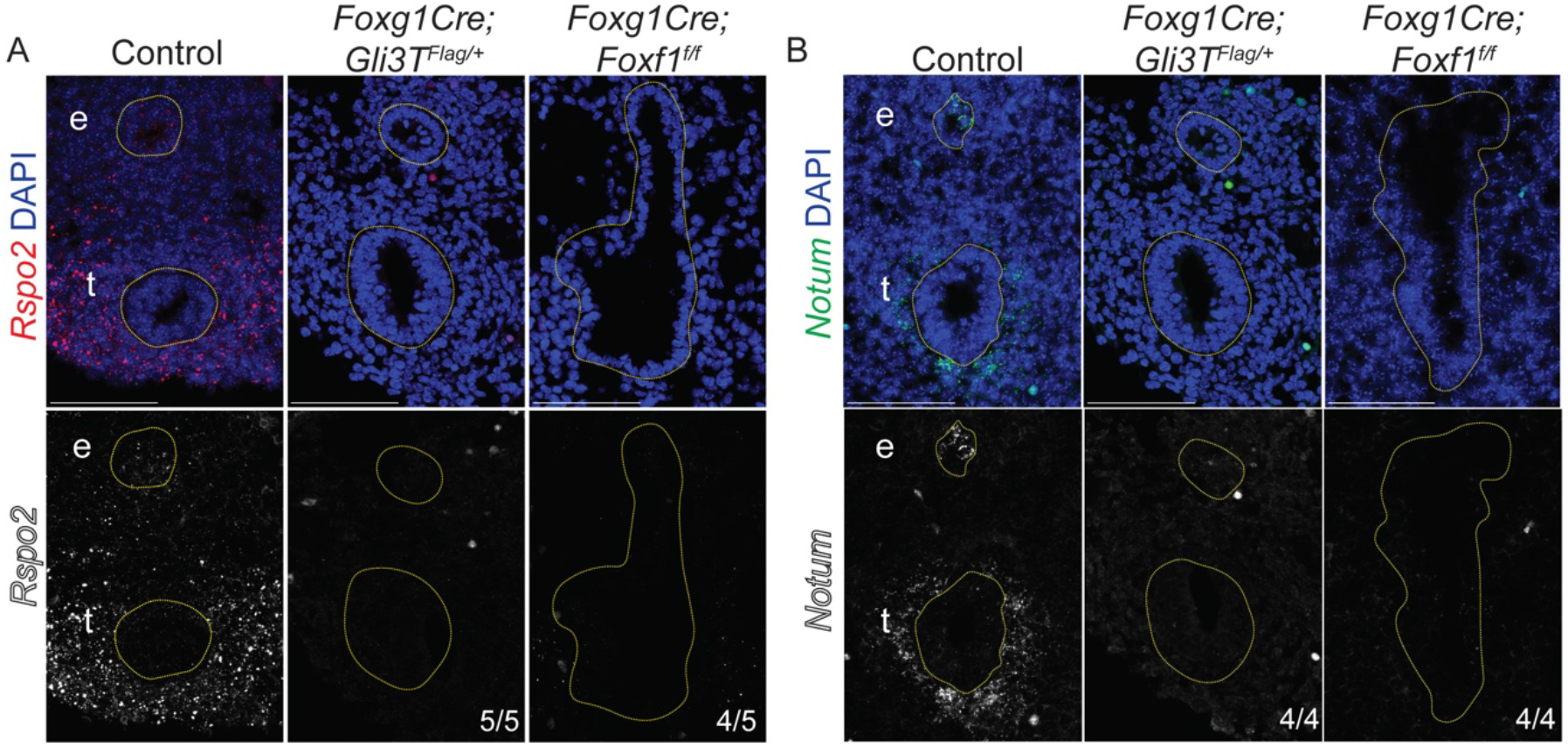
Expression of Wnt pathway genes *Rspo2* and *Notum* are reduced in *Gli* and *Foxf1* Mutants. A-B: RNAscope *in situ* hybridization of E11.5 control, *Foxg1Cre;Gli3T*^*Flag*/+^, and *Foxg1Cre;Foxf1*^*f/f*^ embryos reveals decreases in *Rspo2* (A) and *Notum* (B) in the ventral-lateral tracheal mesenchyme. These results suggest that HH/Gli-Foxf1 signaling is upstream of Rspo2 and Notum during tracheal development. All scale bars, 100 μm. e=esophagus, t=trachea.

## DISCUSSION

In this study we show that conditional mouse mutants with relatively high levels of GliR, mimicking PHS, exhibit tracheomalacia and fail to properly specify Sox9+ tracheal chondrocytes. Our data suggest a model (Figure 7) where: 1) Epithelial HH signaling activate Gli TFs in the mesenchyme to maintain *Foxf1* transcription. 2) Foxf1 in turn is also required for *Sox9* expression at the initiation of tracheal chondrogenesis. 3) Gli3 and Foxf1 TFs act in a regulatory network to promote the transcription of *Sox9, Foxf1*, *Wnt11* and *Notum.* 4) HH/Gli indirectly regulate *Rspo2* and *Wnt4* expression via Foxf1. 5) This Gli-Foxf1-Rspo2 axis promotes Wnt responses in the mesenchyme, which are required for activation of *Sox9* expression and development of the tracheal cartilage (Snowball et al., 2015). The available evidence suggest the Wnt cargo protein Wls is essential for Wnt ligand secretion from epithelium (primarily Wnt7b), which acts together with Rspo2 and other Wnt ligands such as Wnt4 to stimulate *Sox9* expression; mutations in each of these indivudial components results in reduced cartilage (Bell et al., 2008; Caprioli et al., 2015; Gerhardt et al., 2018; Kishimoto et al., 2018; Li et al., 2002; Rajagopal et al., 2008; Snowball et al., 2015). Together, these data provide a mechanistic basis for the tracheomalacia in patients with mutations in HH/Gli pathway genes.

**Figure 7:**
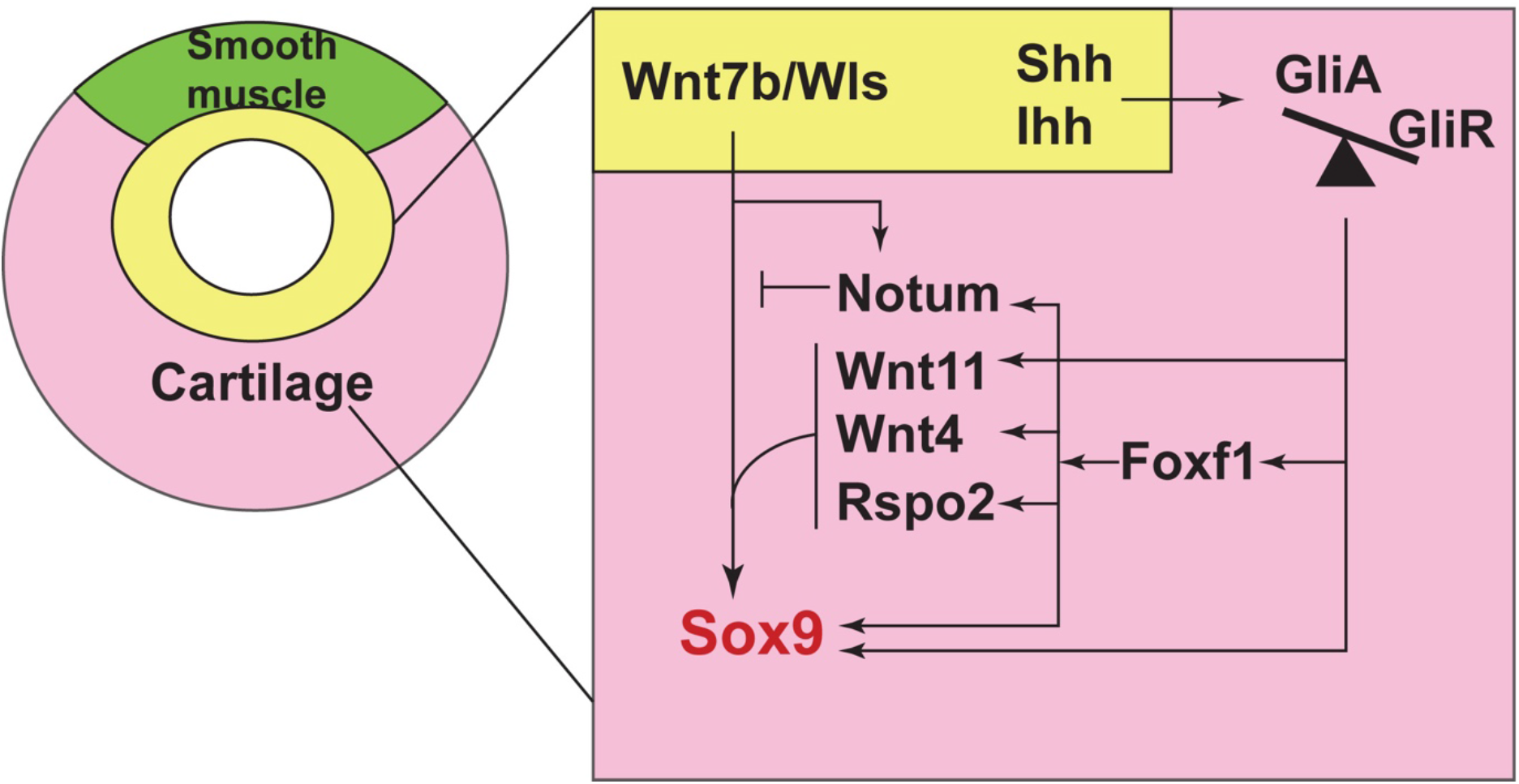
Model of a HH/Gli-Foxf1-Wnt signaling network controlling Sox9+ chondrogenesis. HH/Gli signals from the epithelium (yellow) result in more Gli2/3 activator (GliA) than Gli repressor (GliR). Activated Gli2/3 directly stimulate *Foxf1* during early tracheal development, which supports growth and survival of the tracheal mesoderm (pink). Foxf1 in turn promotes *Sox9* transcription via Wnt/ß-catenin signaling in the ventral tracheal mesoderm to initiate tracheal chondrogenesis. This includes Rspo2, Wnt7b, Wnt5, and Wls, all components of the Wnt signaling pathway that have been previously shown to be required for chondrogenesis. Notum, another direct Gli and Foxf1 target, attenuates Wnt/ß-catenin signaling to regulate maturation of Sox9+ tracheal chondrocytes. In the dorsal tracheal mesenchyme (green), which is thought to have lower Wnt and Bmp signaling, Gli-Foxf1 does not activate Sox9+ chondrogenesis and the tissue adopts a smooth muscle fate.

### Temporal Roles of HH/Gli in Tracheal Chondrogenesis

We took advantage of the different times that *Foxg1Cre* and *Dermo1Cre* transgenes recombine to investigate the temporal-specific role for HH/Gli signaling in tracheal chondrogenesis. The fact that the earlier acting *Foxg1Cre* mutants, which recombines starting at E8.5, had a more severe phenotype that the *Dermo1Cre* mutants, which recombine robustly starting at E10.5, indicates that HH/Gli acts first early in the foregut lateral plate mesoderm at least in part by promoting Foxf1 expression. As previous work has indicated that deletion of *Shh* beginning at E13.5 does not impair tracheal chondrogenesis, the exhibited tracheomalacia in both *Dermo1Cre;Smo*^*f/f*^ and *Dermo1Cre;Gli3T*^*Flag*/+^ mutants suggests a role for HH/Gli signaling between E11.5 and E13.5 in maintaining Foxf1 and Sox9 expression and promoting tracheal chondrogenesis (Miller et al., 2004). Alternatively, Ihh/Gli signaling might compensate for the conditional loss of Shh/Gli activity in conditional *Shh* mutants. We postulate that HH/Gli activity may be acting over a prolonged period of time or that redundant Ihh ligands may compensate for the loss of Shh in that study. Sox9 has previously been established as a direct Gli target gene in other contexts, consistent with our analysis suggesting that a prolonged input of both Gli and Foxf1 may activate and maintain Sox9 expression (Bien-Willner et al., 2007; Tan et al., 2018).

Our study also suggests that the balance of GliA to GliR activity is critical for specification of Sox9+ chondrocytes. The *Smo*^*f/f*^ mutants mimic an absence of HH ligand and GliA function, while *Gli3T*^*Flag*/+^ mutants model human PHS and contain an extra copy of Gli3R (Vokes et al., 2008). Both *Smo*^*f/f*^ and *Gli3T*^*Flag*/+^ mouse models exhibit tracheomalacia and a reduction of Sox9. Previous work showed that *Shh*^−/−^ and *Gli2*^−/−^;*Gli3*^+/−^ mouse mutant models, which have relatively high levels of Gli3R, displayed tracheomalacia, whereas the *Gli2*^+/−^;*Gli3*^−/−^ which lack Gli3R, do not (Litingtung et al., 1998; Miller et al., 2004; Nasr et al., 2019; Park et al., 2010). Previous work suggests that some Hedgehog targets may be activated in the loss of GliR; it is also possible that decreased expression of some HH targets is caused by a reduction in GliA rather than an added presence of GliR (Falkenstein and Vokes, 2014). Future experiments comparing Gli2 and Gli3 transcriptional targets with Gli2 and Gli3 ChIP-seq from the developing foregut at different time points will be important to better define the temporal role of Glis in activating and maintaining the Sox9+ chondrocyte lineage and the relative role of GliA versus GliR.

### Foxf1 Is Required for Specification of Sox9+ Tracheal Chondrocytes

Our analysis indicate that Foxf1 is required for specification of Sox9+ tracheal chondrocytes. Foxf1 is initially expressed throughout the lateral plate mesoderm, but is reduced in the ventral tracheal mesoderm coinciding with the onset of Sox9 activation. We demonstrated with *Foxg1Cre;Foxf1* mutants that Sox9 expression is initially dependent on Foxf1. This result corresponds with the impaired tracheal chondrogenesis and altered tracheal formation observed in *Foxf1* mutants (Mahlapuu et al., 2001; Ustiyan et al., 2018). We postulate that early Foxf1 promotes Sox9 expression in several ways. First, Foxf1 is known to be essential for mesenchymal proliferation and survival in many contexts, and while this alone cannot account for the phenotype we observe, we expect that this contributes to the ultimate expansion of ventral mesenchyme that will ultimately give rise to chondrocytes. In addition, the ChIP-seq data suggests that Foxf1 may directly bind *Sox9* regulatory elements to stimulate its transcription.

However, after E10.5, Foxf1 is downregulated in the ventral mesenchyme as Sox9 expression is upregulated. We largely ruled out the possibility that Foxf1 and Sox9 mutually repress each other’s expression at this time in development. We postulate that feedback regulatory pathways such as Wnt or BMP, both of which are active in the ventral mesoderm, might account for the downregulation of Foxf1, but this remains to be tested (Goss et al., 2009; Li et al., 2008; Snowball et al., 2015).

### HH/Gli Regulates a Foxf1-Rspo2-Wnt Axis

While both HH and Wnt signaling are known to be essential for tracheal chondrogenesis, how these pathways interact was previously unclear. Our analysis indicates that epithelial HH signals stimulate Gli activity in the adjacent ventral mesenchyme to activate a Foxf1-Rspo2-Wnt signaling axis which promotes Sox9 expression. During normal tracheal development, Wnt ligands secreted from the E10.5-13.5 ventral respiratory epithelium are required to signal to the ventral mesenchyme to activate Sox9 expression. Conditional epithelial deletion of the Wnt cargo protein *Wls*, which is required for Wnt ligand secretion, results in a failure to specify Sox9+ chondrocytes (Snowball et al., 2015). A number of other Wnt ligands, including Wnt4, Wnt5a, and Wnt2, are also expressed in the foregut mesenchyme and also contribute to development of the tracheal mesenchyme (Caprioli et al., 2015; Goss et al., 2009; Kishimoto et al., 2018; Li et al., 2002; Snowball et al., 2015). For example, *Wnt4* mutants exhibit reduced and disorganized tracheal cartilage (Caprioli et al., 2015).

Our data indicate that Gli and Foxf1 promote the expression of Wnt4, Rspo2 and Notum in the tracheal mesenchyme. Rspo2 is a secreted ligand that binds Lgr/Lrp6 receptor complexes to potentiate canonical Wnt signaling, while Notum is a Wnt target and acts in a negative feedback loop to inhibit Wnt signaling (Bell et al., 2008; Gerhardt et al., 2018). Both Rspo2 and Notum are required for normal tracheal chondrogenesis: *Rspo2* mutant tracheas show a reduction in the number of tracheal cartilage rings, while *Notum* mutant tracheas demonstrate impaired tracheal cartilage differentiation (Bell et al., 2008; Gerhardt et al., 2018). Importantly, our data indicate that *Rspo2*, *Wnt4*, *Wnt11*, and *Notum* appear to be direct targets of Foxf1 in the developing respiratory system, suggesting that HH/Gli signaling may act through Foxf1 to support Wnt activity through these Wnt pathway members during tracheal chondrogenesis. Previous reports indicate that Gli acts in a combinatorial fashion to directly upregulate Foxf1 and chondrogenic targets, suggesting that Gli and Foxf1 might act together to mediate expression of *Sox9* and *Notum* in the context of tracheal development (Hoffmann et al., 2014; Tan et al., 2018). ChIP sequencing of Gli3 and Foxf1 performed in the E10.5 and E11.5 foregut would more directly show that these four Wnt signaling components are directly downstream of HH/Gli-Foxf1.

Together, loss of *Rspo2, Wnt4* and *Notum* may account for the failure in activation of Sox9 expression in *Foxg1Cre* mutants. As tracheomalacia observed in *Rspo2* mutants is made worse with additional reduction in *Lrp6*, this suggests that a combinatorial disruption in Wnt activity may severely impact normal tracheal chondrogenesis (Bell et al., 2008). Indeed, Sox9 expression may be absent from the dorsal tracheal mesenchyme since few Wnt and BMP signaling members are presented in the dorsal tracheal mesenchyme to support HH/Gli-mediated Sox9+ cartilage specification there (Gerhardt et al., 2018; Li et al., 2008; Snowball et al., 2015). While Wnt-ß-catenin signaling appears to repress *Sox9* in the context of limb chondrogenesis, *Sox9* is dependent on Wnt-ß-catenin during gut development, indicating that Wnt regulation of Sox9 may be context-dependent (Blache et al., 2004; Kozhemyakina et al., 2015). Gli1 has also been shown to directly bind to a *Sox9* enhancer and support *Sox9* expression (Bien-Willner et al., 2007). Thus, it is also possible that the Wnt effector transcription factors such as Tcf proteins and ß-catenin, once induced by Rspo2-Lrp6 interaction, co-regulate expression of Sox9 with Gli and Foxf1. Altogether, our data provides potential mechanisms through which disruptions in HH/Gli signaling may impair specification of Sox9+ tracheal chondrocytes, ultimately leading to tracheomalacia.

## MATERIALS AND METHODS

### Animal models

All mouse experiments were approved by the Institutional Animal Care and Use Committee at Cincinnati Children’s Hospital Medical Center (CCHMC) under protocol 2019-0006. Animals were housed within the CCHMC Veterinary Services Core in temperature-controlled rooms with regular access to food and water. Dr. Debora Sinner provided *Foxg1Cre* (Hebert and McConnell, 2000), *Dermo1Cre* (Sosic et al., 2003), and *mTmG* (Muzumdar et al., 2007) animals, as well as *Foxg1Cre;Sox9*^*f/f*^ (Kist et al., 2002) mutant and control samples. Dr. Vladimir Kalinichenko provided *Foxf1*^*fl/fl*^ animals (Ren et al., 2014). Dr. Samantha Brugmann provided *Wnt1Cre* (Lewis et al., 2013)*, mTmG* (Muzumdar et al., 2007), and *Gli3T*^*Flag/Flag*^ (Vokes et al., 2008) animals. Dr. Joo-Seop Park provided *Gli3T*^*Flag/Flag*^ animals (Vokes et al., 2008).

### Immunostaining, *in situ* hybridization, and Alcian Blue staining

At least three embryos of each genotype were used for all experiments. For section immunostaining, embryos were collected and incubated in 4% paraformaldehyde solution overnight at 4°C. After two rinses in 1XPBS, embryos were incubated in 30% sucrose overnight at 4°C before embedding in OCT for cryosectioning. Sections were collected at 8 μm. *Foxg1Cre;Sox9*^*f/f*^ embryos were embedded in paraffin before sectioning, and were deparaffinized before beginning immunostaining. On the first day of immunostaining, sections were washed in 1XPBS before incubation in 1XPBS with 0.05% Triton X-100. Sections were then blocked with 5% normal donkey serum in 1X PBS for one hour before overnight incubation in primary antibodies (Foxf1, goat, R&D, 1:300; Sox9, mouse, Invitrogen, 1:200; Sox9, rabbit, Millipore, 1:200; Acta2, mouse, Sigma, 1:800; Acta2, rabbit, Genetex, 1:800; pHH3, mouse, Millipore, 1:1000; CC3, rabbit, Cell Signaling, 1:200; GFP, chicken, Aviva Biosystems, 1:1000; DsRed, mouse, Living Color, 1:1000, ß-galactosidase/LacZ, chicken, Abcam, 1:1000) at 4°C. On the second day, sections were washed three times in 1XPBS before incubation in secondary antibodies (donkey anti-mouse 647, Jackson; donkey anti-goat 647, Jackson; donkey anti-rabbit 647, AlexaFluor; donkey anti-rat 647, Jackson; donkey anti-rabbit Cy3, Jackson; donkey anti-mouse Cy3, Jackson; donkey anti-goat 488, Jackson; donkey anti-chicken 488, Jackson; donkey anti-chicken 647, Jackson; donkey anti-rabbit 405, Abcam; DAPI, ThermoScientific, all at 1:500) at room temperature, and were washed in 1XPBS three more times before coverslip placement.

*In situ* hybridization was performed using an RNAScope Multiplex Fluorescent v2 kit according to manufacturer instructions (ACD Biosystems). For wholemount immunostaining, embryos were stored in methanol at −20°C before beginning staining. Foreguts were dissected out, incubated in Dent’s Bleach for two hours, and were serially rehydrated into 1XPBS before blocking in 5% normal donkey serum and 1% DMSO for two hours. Foreguts were then incubated in primary antibody diluted in blocking solution overnight at 4°C. After five washes in 1XPBS, foreguts were incubated in secondary antibody overnight at 4°C. The next day, after three washes in 1XPBS, foreguts were serially dehydrated into methanol and stored at 4°C overnight before clearing in Murray’s Clear solution for imaging. All images were taken on a Nikon LUNA upright confocal microscope in the CCHMC Confocal Imaging Core. All scale bars are 100 μm.

Alcian Blue staining was performed on dissected foreguts as previously described (Que et al., 2007). Foreguts were then serially rehydrated into 1XPBS before incubation in 30% sucrose overnight at 4°C. After embedding in OCT, foreguts were cryosectioned at 60 μm thickness and photographed using a Nikon LUN-A inverted widefield microscope.

### Quantitative analysi

Confocal images were analyzed using Nikon Elements Analysis and Imaris programs. All statistical analyses were performed in Microsoft Excel on data obtained from single transverse sections from each embryo. Sections were selected based on their median location between the anterior separation of the larynx into the trachea and the posterior formation of the mainstem bronchi from the trachea. Calculations were performed in Microsoft Excel using a two-sided Student’s t-test with unequal variance and with significance defined as p<0.05. Graphs were generated using Prism. For relative expression of Foxf1 and Sox9 in the tracheal mesoderm (referred to as TMes in figures), the number of Foxf1+ or Sox9+ tracheal mesoderm cells was divided by the total number of tracheal mesoderm cells. For pHH3+ (mitotic indices) or CC3+ rates of mesenchyme cells, the number of tracheal mesenchymal cells positive for either pHH3 or CC3 was divided by the total number of tracheal mesenchymal cells.

### RNA-seq and ChIP-seq analysis

RNA-Seq analysis was performed on control and *Foxg1Cre;Gli3T*^*Flag*/+^ samples sequenced at stages E10.5 (foreguts) and E11.5 (tracheas) to study tracheal congenital defects. After storing in RNALater (Ambion) at −80°C, RNA was isolated using a Qiagen MicroEasy Kit and was amplified by the CCHMC Gene Expression Core before sequencing in the CCHMC DNA Sequencing Core using an Illumina 3000 high throughput platform. Single end sequencing with read-depth of ~22-27 million and read length of 75bp was performed. Raw reads from the experiments were analyzed using Computational Suite for Bioinformaticians and Biologists (CSBB −v3.0, https://github.com/praneet1988/Computational-Suite-For-Bioinformaticians-and-Biologists). The following steps were carried out in analysis using CSBB. Quality check and trimming was performed using FASTQC (https://www.bioinformatics.babraham.ac.uk/projects/fastqc/) and Bbduck (https://jgi.doe.gov/data-and-tools/bbtools/bb-tools-user-guide/bbduk-guide/) respectively. Quality trimmed reads were then mapped to the mouse genome (mm10) using Bowtie2 and quantified using RSEM (https://bmcbioinformatics.biomedcentral.com/articles/10.1186/1471-2105-12-323). Differential expression analysis was carried out using CSBB-Shiny (https://github.com/praneet1988/CSBB-Shiny) and volcano plots were generated using EnhancedVolcanco (https://www.bioconductor.org/packages/release/bioc/vignettes/EnhancedVolcano/inst/doc/EnhancedVolcano.html) package in R. Differentially expressed genes were obtained at following thresholds: LogFC > |1| and 0.05 > False Discovery Rate (FDR). ChIP-Seq analysis was performed on following published datasets: 1) Foxf1 ChIP on dissected E18.5 lung, GSE77159 (Dharmadhikari et al., 2016); 2) Gli3-3xFlag ChIP on dissected E10.5 limb buds, GSE133710 (Lex et al., 2020); 3) ATAC-seq and H3K4me3 ChIP performed on E9.5 cardiopulmonary progenitors, GSE119885 (Steimle et al., 2018). These datasets were reprocessed using CSBB and for visualization purpose bigwig files were generated using deeptools (BamCoverage function) (Ramirez et al., 2016) from bam files. Peaks were called using Macs2 (default parameters) (Zhang et al., 2008). Genome browser views were generated using IGV.

## ACKNOWLEDGEMENTS

We would like to thank members of the Zorn and Wells labs as well as Samantha Brugmann, Sheila Bell, and Steve Vokes for critical feedback. We also thank Leslie Brown, Kaulini Burra, and Natalia Bottasso Arias for technical assistance. We are grateful to the staff of the CCHMC Confocal Imaging Core, Veterinary Services Core, Gene Expression Core, and DNA Sequencing Core for their assistance.

## DATA AVAILABILITY

The RNA-seq data has been deposited into GEO: GSE Accession # pending.

## CONTRIBUTIONS

T.N. and A.M.Z. designed experiments and wrote the manuscript. T.N., P.C., and K.A. performed bioinformatics analysis. T.N., J.L.K., K.D., V.U., J.M.S., S.L.T., and D.S. performed experiments. V.V.K., J.M.W., D.S. and A.M.Z. provided support.

## SUPPLEMENTAL FIGURES AND FIGURE LEGENDS

**Supplemental Figure 1:**
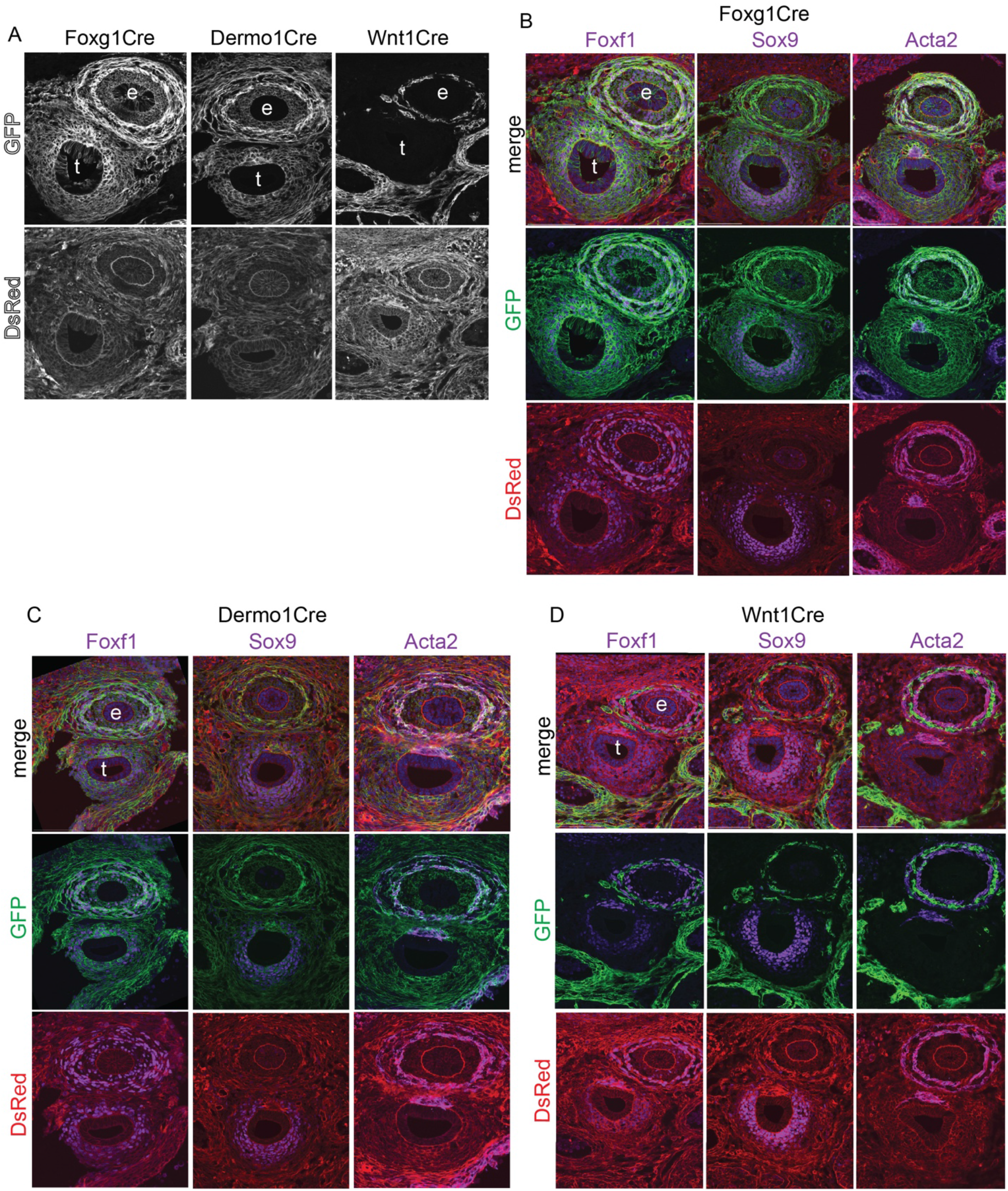
Foxg1, Dermo1, and Wnt1 Lineages during Tracheoesophageal Development. A: GFP and DsRed immunostaining of E13.5 *Foxg1Cre;mTmG*, *Dermo1Cre;mTmG*, and *Wnt1Cre;mTmG* foreguts show that Foxg1 and Dermo1 lineages give rise to tracheoesophageal mesenchyme, while the Wnt1 neural crest cell lineage gives rise only to neurons that appear to innervate the esophageal and trachealis muscles. Foxg1-derived cells can also be found in the endoderm of both the trachea and esophagus. B: GFP and DsRed immunostaining in E13.5 *Foxg1Cre;mTmG* embryos also stained with Foxf1, Sox9, or Acta2. The Foxg1 lineage includes Foxf1+ tracheoesophageal mesoderm, Sox9+ tracheal chondrocytes, and Acta2+ esophageal and tracheal muscles. C: GFP and DsRed immunostaining in E13.5 *Dermo1Cre;mTmG* embryos also stained with Foxf1, Sox9, or Acta2. The Dermo1 lineage can also be found through the tracheoesophageal mesoderm, including smooth muscle and chondrocytes. D: GFP and DsRed immunostaining in E13.5 *Wnt1Cre;mTmG* embryos also stained with Foxf1, Sox9, or Acta2. The Wnt1 lineage appears to give rise to neurons that innervate the esophageal and tracheal smooth muscle, but does not include tracheoesophageal mesoderm. All scale bars, 100 μm. e=esophagus, t=trachea.

**Supplemental Figure 2:**
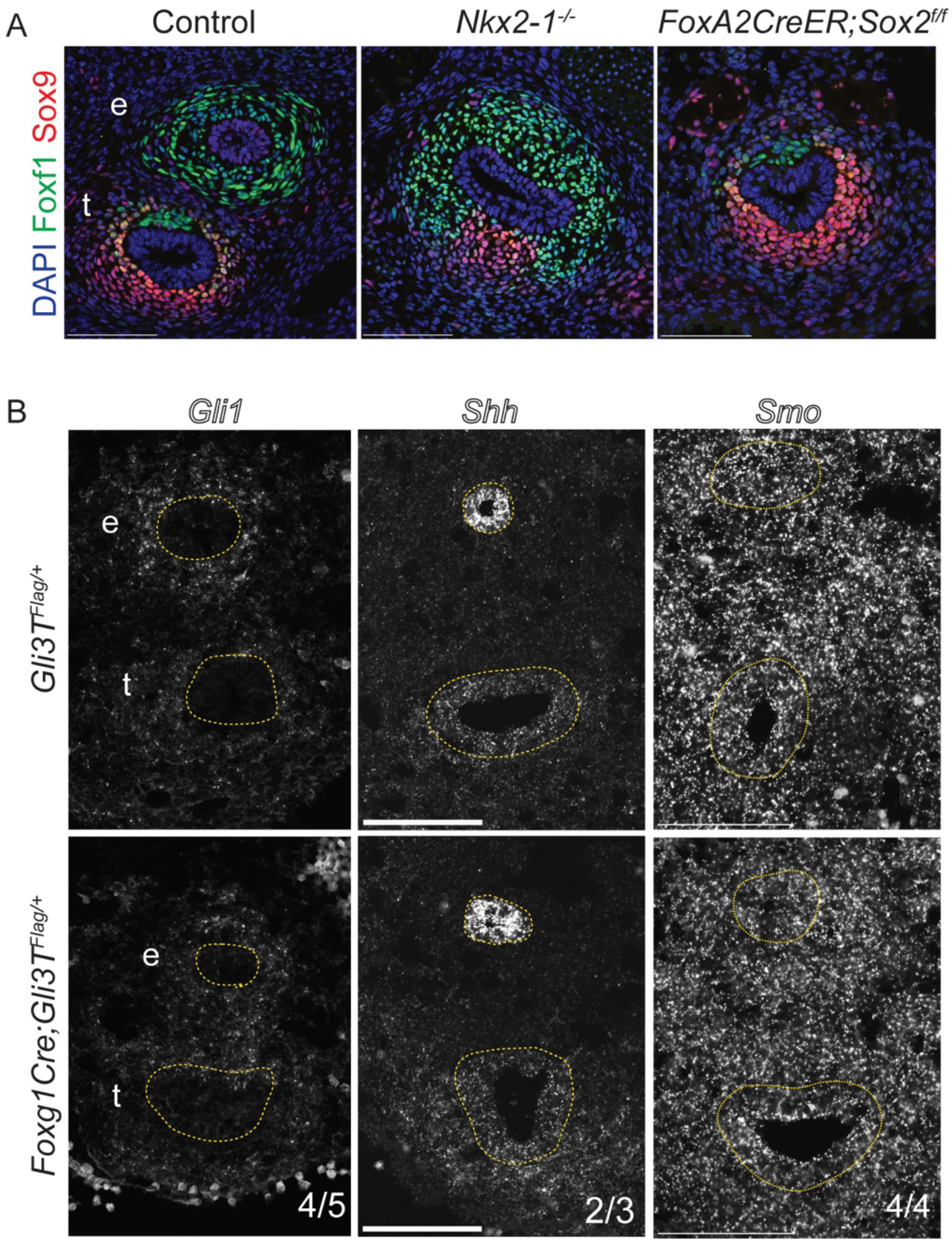
Foxf1 and Sox9 Expression in *Sox2* **and***Nkx2-1* Mutants. A: Immunostaining of Foxf1 and Sox9 in E13.5 *FoxA2CreER;Sox2*^*f/f*^ and *Nkx2-1*^*GFP/GFP*^ (or *Nkx2-1*^−/−^) mutants. *Sox2* mutants exhibit dorsal-ventral patterning of Foxf1 and Sox9 similar to that of the control trachea. *Nkx2-1* mutants exhibit dorsal expansion of Foxf1 but preserved ventral upregulation Sox9 and downregulation of Foxf1. B: RNAscope *in situ* hybridization of *Gli1*, *Shh*, and *Smo* in E11.5 control and *Foxg1Cre;Gli3T*^*Flag*/+^ tracheas suggest that HH/Gli signaling is not significantly deterred by the presence of an extra copy of Gli3 repressor. All scale bars, 100 μm. e=esophagus, t=trachea.

**Supplemental Figure 3:**
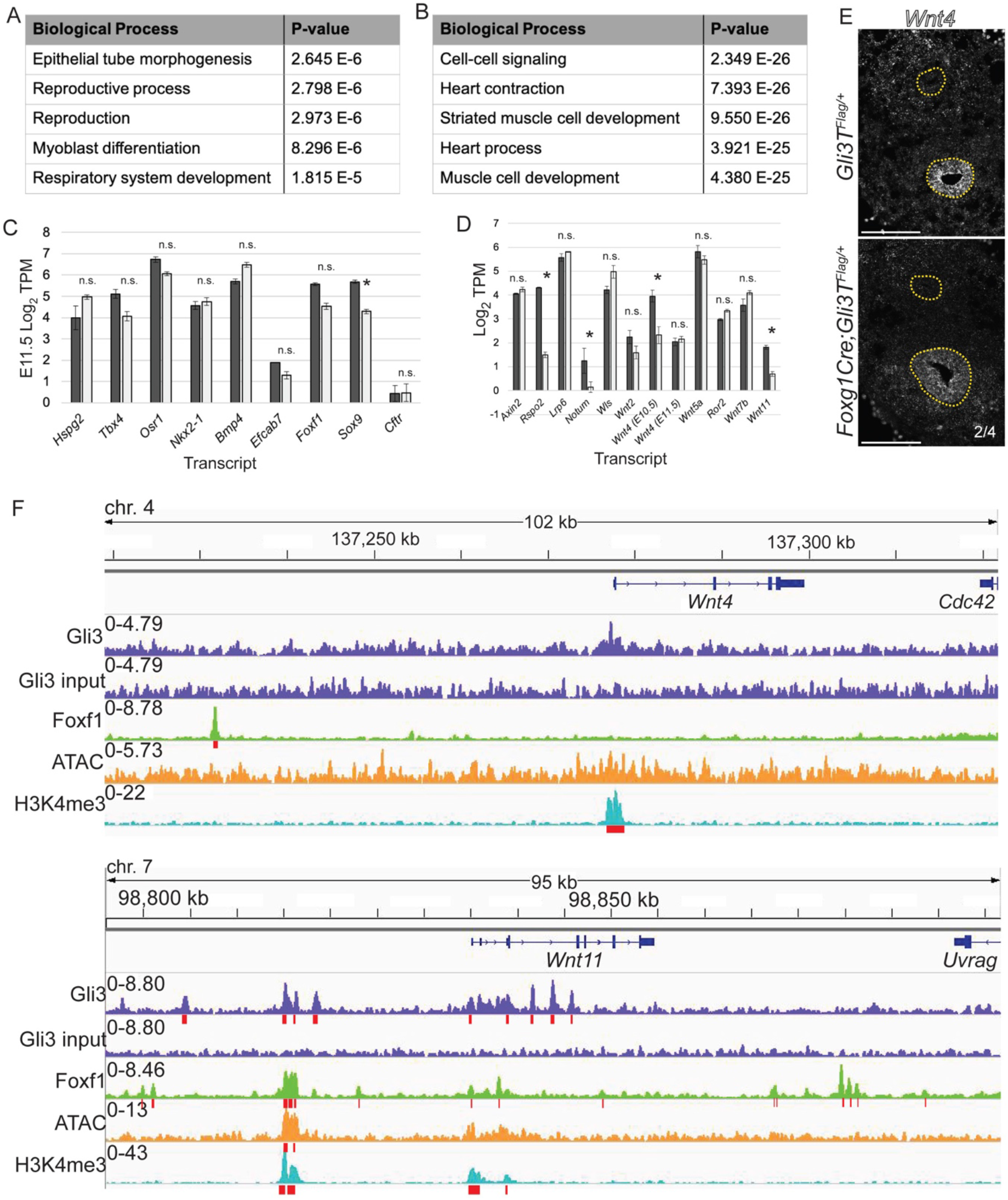
Expression of Select Mediators of Tracheal Mesenchymal Development in *Foxg1Cre;Gli3T*^*Flag*/+^ Mutants. A-B: GO enrichment analysis for downregulated genes in *Foxg1Cre;Gli3T*^*Flag*/+^ mutants (E10.5 and E11.5 combined) and B) upregulated genes. C: Histogram of RNA-seq transcript levels of selected known mediators of tracheal chondrogenesis (Sinner et al., 2019). Asterisks indicate differential expression of Log_2_TPM≥|1|, p<0.05 in E11.5 *Foxg1Gli3T*^*Flag*/+^ tracheas. D: Histogram of RNAseq transcript levels of Wnt signaling pathway members. Asterisks indicate differential expression of Log_2_TPM≥|1|, p<0.05 in E11.5 *Foxg1Gli3T*^*Flag*/+^ tracheas and E10.5 *Foxg1Gli3T*^*Flag*/+^ foreguts (for E10.5 *Wnt4* only). E: Genome browser views of Gli3-3xFlag (GSE133710), Foxf1 (GSE77159), and H3K4me3 (GSE119885) ChiP-seq data, as well as ATAC-seq data (GSE119885) in the *Wnt4* and *Wnt11* loci. *Wnt4* appears to be a direct Foxf1 target, while *Wnt11* may be directly regulated by both Gli3 and Foxf1 (Ustiyan et al., 2018). F: RNAscope *in situ* hybridization of *Wnt4* in E11.5 control and *Foxg1Cre;Gli3T*^*Flag*/+^ tracheas.

